# G-quadruplex stabilization in the ions and maltose transporters inhibit *Salmonella enterica* growth and virulence

**DOI:** 10.1101/357046

**Authors:** Neha Jain, Subodh Kumar Mishra, Uma Shankar, Arpita Tawani, Ankit Jaiswal, Tarun Kumar Sharma, Prashant Kodgire, Amit Kumar

**Affiliations:** Discipline of Biosciences and Biomedical Engineering, Indian Institute of Technology Indore, Indore, Simrol, Indore, 453552, India; Centre for Bio-design and Diagnostics, Translational Health Science and Technology Institute, Faridabad, Haryana, India.

## Abstract

The G-quadruplex structure forming motifs have recently emerged as a novel therapeutic drug target in various human pathogens. Herein, we report three highly conserved G-quadruplex motifs (SE-PGQ-1, 2, and3) in genome of all the 412 strains of *Salmonella enterica*. Bioinformatics analysis inferred the presence of SE-PGQ-1 in the regulatory region of *mgtA*, presence of SE-PGQ-2 in the open reading frame of *entA* and presence of SE-PGQ-3 in the promoter region of *malE* and *malK* genes. The products of *mgtA* and *entA* are involved in transport and homeostasis of Mg^2+^ and Fe^3+^ ion and thereby required for bacterial survival in the presence of reactive nitrogen/oxygen species produced by the host macrophages, whereas, *malK* and *malE* genes are involved in transport of maltose sugar, that is one of the major carbon source in the gastrointestinal tract of human. The formation of stable intramolecular G-quadruplex structures by SE-PGQs was confirmed by employing CD, EMSA and NMR spectroscopy. Cellular studies revealed the inhibitory effect of 9-amino acridine on *Salmonella enterica* growth. Next, CD melting analysis demonstrated the stabilizing effect of 9-amino acridine on SE-PGQs. Further, polymerase inhibition and RT-qPCR assays emphasize the biological relevance of predicted G-quadruplex in the expression of PGQ possessing genes and demonstrate the G-quadruplexes as a potential drug target for the devolping novel therapeutics for combating *Salmonella enterica infection*.

**Author Summary:** Since last several decades’ scientific community has witnessed a rapid increase in number of such human pathogenic bacterial species that acquired resistant to multiple antibacterial agents. Currently, emergence of multidrug-resistant strains remain a major public health concern for clinical investigators that rings a global alarm to search for novel and highly conserved drug targets. Recently, G-quadruplex structure forming nucleic acid sequences were endorsed as highly conserved Drug target for preventing infection of several human pathogens including viral and protozoan species. Therefore, here we explored the presence G-quadruplex forming motif in genome of *Salmonella enterica* bacteria that causes food poisoning, and enteric fever in human. The formation of intra molecular G-quadruplex structure in four genes (*mgtA*, *entA*, *malE* and *malK*) was confirmed by NMR, CD and EMSA. The 9-amino acridine, a known G-quadruplex binder has been shown to stabilize the predicted G-quadruplex motif and decreases the expressioin of G-quadruplex hourbouring genes using RT-PCR and cellular toxicity assay. This study concludes the presence of G-quadruplex motifs in essential genes of *Salmonella enterica* genome as a novel and conserved drug target and 9-amino acridine as candidate small molecule for preventing the infection of *Salmonella enterica* using a G4 mediated inhibition mechanism.

## Introduction

*Salmonella enterica* belongs to *Enterobacteriaceae family* and known to cause typhoid fever and food poisoning in the human. *Salmonella enterica* consists of six subspecies namely 1) *enterica* 2) *salamae* 3) *arizonae* 4) *diarizonae* 5) *houtenae* 6) *indica* (Fig 1a). Among these subspecies, subspecies *S. enterica* is known to infect human and possess ~*2463 serovars* that can be further divided into two subclasses[1, 2]. Typhoidal class of *S. enterica* included *S. enterica subsp. enterica ser*. Typhi(S. ser. Typhi) and *S. enterica subsp. enterica ser*. Paratyphi (*S*. ser. Paratyphi) known to causes typhoid or enteric fever, whereas non-typhoidal class includes S. *enterica subsp. enterica ser*. Enteritidis (*S*. ser. Enteritidis) and *S. enterica subsp. enterica ser*. Typhimurium (*S*. ser. Typhimurium), and causes food poisoning in human[3]. They have been reported for causing more than ~140 foodborne illness throughout the world [4–8]. As per the Centers for Disease Control and Prevention (CDC), typhoid fever causes ~22 million new cases and ~200,000 deaths every year across the world [9, 10]. The emergence of antimicrobial drug resistance for chloramphenicol, co-trimoxazole, ampicillin, ciprofloxacin, ofloxacin, azithromycin and cephalosporin make the situation more dangerous and thus leading to increased death rate due to clinical treatment failure [11–13]. More recently, severe complications with *Salmonella* species have emerged and connected to multiple invasive disorders like irritable bowel syndrome [14], reactive arthritis[15], bacteremia[16] focal infection[17], meningitis[18] and infectious aortitis[19]. Due to its high prevalence and rapid emergence of drug resistance, *Salmonella* ring a global alarm for the development of novel and promising therapeutic approaches. A more effective and efficient therapeutic approach would be required to target the expression of those genes that were previously observed to be associated with essential nutrient acquisition system and remained conserved throughout the evolution in the genome of this deadly human pathogen.

**Fig. 1.**
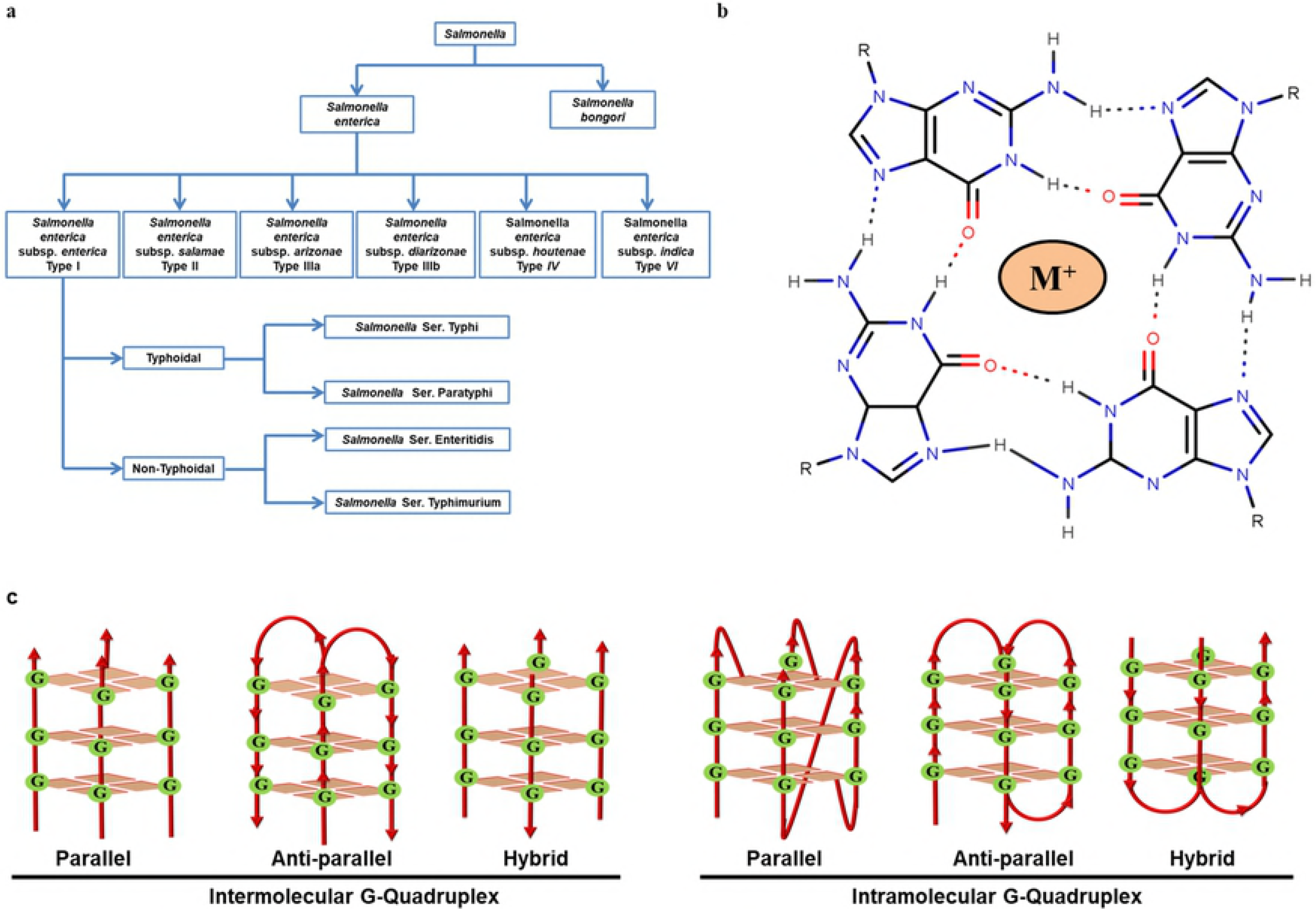
*Salmonella* genus classification and G-quadruplex topologies. a) Flowchart depicting the classification of the *Salmonella* genus. **b)** G-quartet structure showing hoogsteen hydrogen bond formation between Guanines and cation binding. **c)** Different types of topologies formed by intermolecular and intramolecular G-quadruplex structures.

*S. enterica* is an intracellular pathogen that grows in phagocytes and macrophages. During the growth phase, the host innate immune system generate various oxidative stresses to eradicate this pathogen. However, *S. enterica* possess a magnesium homeostasis mechanism that controls the intracellular Mg^2+^ concentration and helps bacteria to survive in nitro-oxidative stressed condition[20]. Magnesium homeostasis is also required for virulence, thermo-tolerance and survival of S. *enterica [21]*. The bacteria contain three genes that help in Mg^2+^ uptake from the host namely: i) *mgtA*, ii) *mgtB*, and iii) *corA*. Neutralization of the reactive nitrogen stress(RNS, nitro-oxidative) is mainly regulated by Mg^2+^ transport ATPase that is encoded by *mgtA* gene and therefore plays a vital role in the bacterial survival inside macrophage (Fig 2a) [22]. Hence, targeting the conserved region of *mgtA* gene may serve as a promising therapeutic approach to combat with *S. enterica* pathogenesis.

**Fig. 2.**
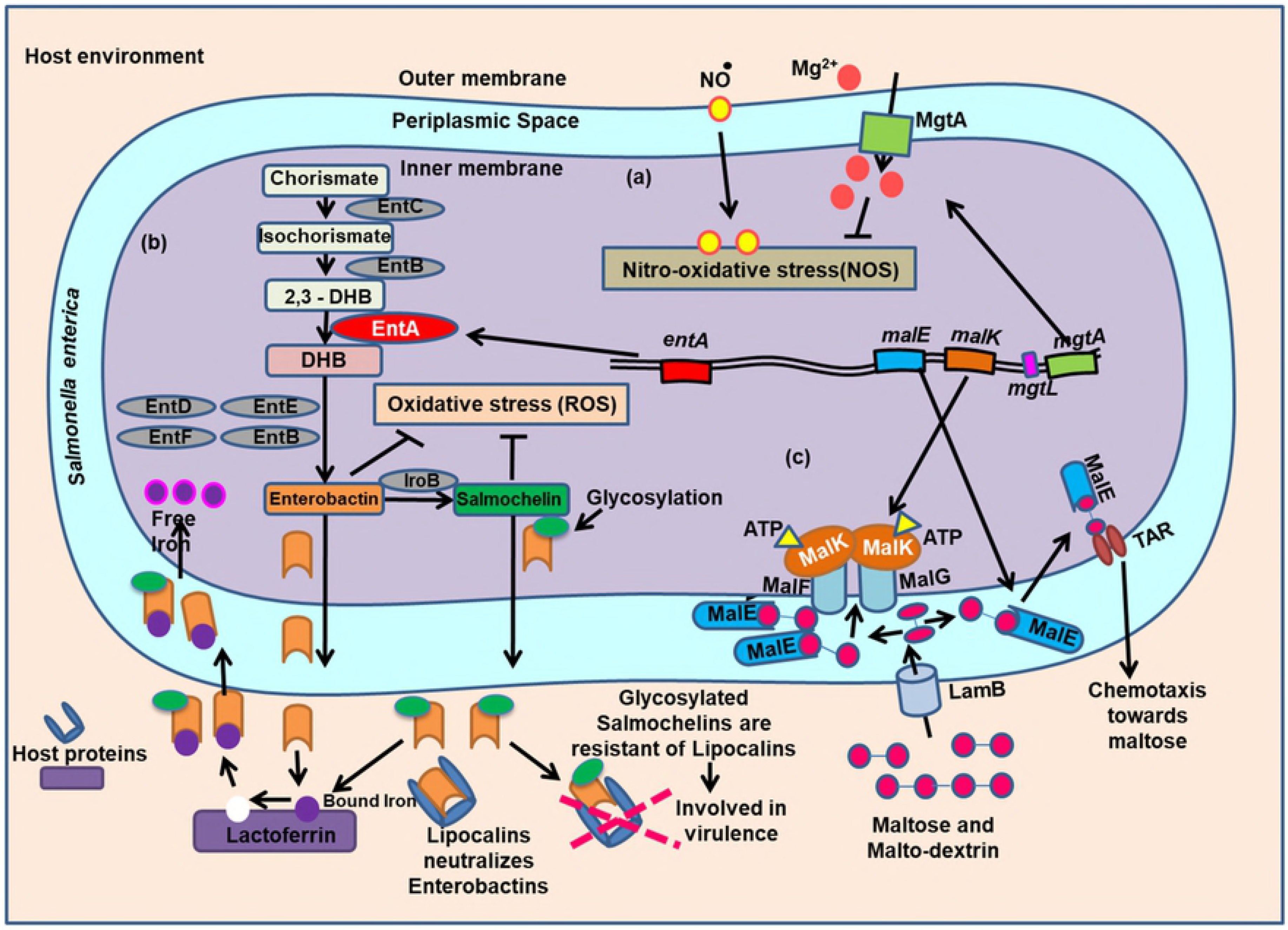
*mgtA*, *entA* and maltose operon mediated mechanisms. Schematic representation of the functions related with a) *mgtA*. b) *entA* and c) maltose operon mediated mechanisms in growth and virulence of *Salmonella enterica*

Similar to Mg^2+^ ion, iron is also required by this bacteria, but its uptake from the host environment is more challenging as it gets sequestered by various host proteins like siderocalin, transferrin and lactoferrin leading to its unavailability in the host environment (Fig 2b). *S. enterica* produces enterobactin and salmochelin, two low molecular weight catecholate type siderophores that have a high affinity for Iron than the host iron-binding proteins. These enterobactin/salmochelin are produced from chorismate that requires several enzymes encoded by a co-transcribed *ent*ABCDEF operon (Fig 2b). [23]. The conversion of the 2,3-dihydro-2,3-dihydroxybenzoate (2,3 DHB) to another intermediate 2,3-dihydroxybenzoate(DHB) is a crucial step of this pathway and required a 2,3-dihydro-2,3-dihydroxybenzoate dehydrogenase enzyme(EntA), encoded by *entA* gene of *ent*ABCDEF operon. The inhibition of *entA* gene expression has been observed to abolish the DHB formation leading to the reduced production of bacteriocin and salmochelin[24]. Interestingly, salmochelin are resistant to antimicrobial peptides (lipocalins), secreted by the host cells and acts as an essential factor in the pathogenesis of systemic S. ser.Thyphimurium infection [25]. Salmoechelins also protects this bacteria from reactive oxygen superoxides (ROS) mediated oxidative stress[26]. Targeting the production of siderophore has previously shown to have antimicrobial activity against *Mycobacterium tuberculosis* [27], *Aspergillus fumigatus*[28], *Yersinia pestis*[29], *Pseudomonas aeruginosa*[30], *Bacillus substilis*, *Acinetobacter baunamnni* and *Vibrio chloreae*[31]. Therefore, targeting *entA* gene may prove as another therapeutic approach to combat the infection and virulence of *S. enterica* as well.

*S. enterica* grows in the gastrointestinal tract of humans that has an ample amount of maltose and maltodextrin. Therefore, along with glucose, *S. enterica* utilizes maltose as major source of carbon. The uptake of maltose from host environment is tightly regulated by two genes, *malK* and *malE*(Fig 2c) [32]. Inhibiting the *malK* and *malE* synthesis have shown to decline the growth rate of *S. enterica* [33]. Henceforth, targeting these genes will make *S. enterica* unable to grow inside the gastrointestinal tract.

DNA along with its canonical B-form can also fold into a non-canonical G-quadruplex (G4) structure. G rich regions having a motif G_2_N_*L*_G_≥2_N_*L*_G_≥2_N_*L*_G_≥2_ present in the genome tends to form specific secondary structures known as G-quadruplex (G4). G4s are stabilized by the presence of monovalent and some divalent cations, in the order of K^+^ > Na^+^ > Mg^2+^ > Li^+^ and can adopt various topologies (Fig 1b & 1c) [34, 35]. This structural diversity has been exploited for the diagnosis and therapeutic targeting[36, 37]. G4s are highly ordered and shown to be evolutionarily conserved in eukaryotes [38] prokaryotes[39], protista[40], plants[41] and viruses[42]. Presence of G4 binding proteins and antibody-based approaches have confirmed their presence *in-vivo* and are reported to play a regulatory role in the expression of genes such as regulating DNA replication by the specification of origin of replication(ORI) sites, telomere maintenance in human cells, antigenic variations by regulating recombination, transcription, and translation[43]. G4 motifs present in the telomere regions of the chromosomes and promoter regions of oncogenes like Bcl, c-Kit, MYC, KRAS, etc. have been explored in various human cancers[44, 45].

Currently, G4s are being investigated for their involvement in virulence and survival mechanisms of various human pathogen[46]. Stabilization of G4s in protozoans: *Plasmodium falciparum*, *Trypanosoma brucei* and *Leishmania donovani* have shown anti-protozoal activities [47]. Further, G4s have emerged as promising drug target in viruses like SARS coronavirus, Human Papilloma virus(HPV), Zika virus, Ebola virus, Herpes simplex virus(HSV), Epstein-Barr virus(EBV), Hepatitis B virus(HBV), Hepatitis C virus(HCV), Human immunodeficiency virus 1(HIV-1), etc. where they play an essential role in viral proliferation and pathogenicity[42]. In bacteria, G4 present at the upstream of *pilE* locus, *B31 vlsE* locus and *tprK* antigen protein in *Neisseria gonorrhoea*, *Borrelia burgdorferi*, and *Treponema pallidum*, respectively acts as an activator for the initiation of antigenic variation and helps the pathogens in bypassing immune system of the host cells[46]. In *Deinococcus radiodurans*, G4 sequences were present in the regulatory regions of various genes and contributes to radio resistance [48]. G4 present at the 150 nt upstream of *nasT* in the soil bacterium *Paracoccus denitrificans PD1222* isreported to be involved in nitrite assimilation[49].

All these reports demonstrated the pivotal role of G-quadruplex in human pathogens and their conserved-ness suggest them as a promising drug target for both drug susceptible and drug-resistant strains of human pathogens. Therefore, a comprehensive study that discovers highly conserved G-quadruplex in the genome of *S. enterica* as a drug target may provide as a most suitable therapeutic approach for fighting against the infection of this deadly pathogen and overcome the emergence of drug-resistant problem in this bacterium.

In present study, we sought to explore the highly conserved potential G-quadruplex forming sequences (SE-PGQs) in all the available and completely sequenced 412 strains of *S. enterica*. Bioinformatics sequence analysis revealed the presence of three SE-PGQs (SE-PGQ 1-3) in three different gene locations of *S. enterica* genome. The SE-PGQ-1 was found to be present in the regulatory region of *mgtA*, SE-PGQ-2 in the open reading frame of *entA*, whereas SE-PGQ-3 found to lie in the regulatory region of *malK* and *malE* genes (Fig 3a). In order to confirm the formation of G-quadruplex structure by SE-PGQs, Circular Dichroism spectroscopy (CD), gel mobility shift assay (EMSA) and one dimensional ^1^H NMR spectroscopy were employed. Further to validate these SE-PGQ as a potential drug target, CD melting, and Taq polymerase stop assay were performed that confirmed 9-amino-acridine, a known G4 binding molecule interact and stabilizes the SE-PGQs motifs with high affinity and selectivity over the duplex DNA. Disc diffusion assay and MTT cytotoxicity assay confirmed the growth inhibition of *S. enterica* cells by 9-amino acridine molecule. Further, Real-time-quantitative PCR (RT-qPCR) revealed the reduced expression of genes that harbor the SE-PGQs in either their coding region or regulatory region upon the treatment with 9-amino acridine. This change in the expression of PGQs harboring gene in the presence of G4 binding ligand suggested a G4 mediated regulatory mechanism in the expression of these genes.

**Fig. 3.**
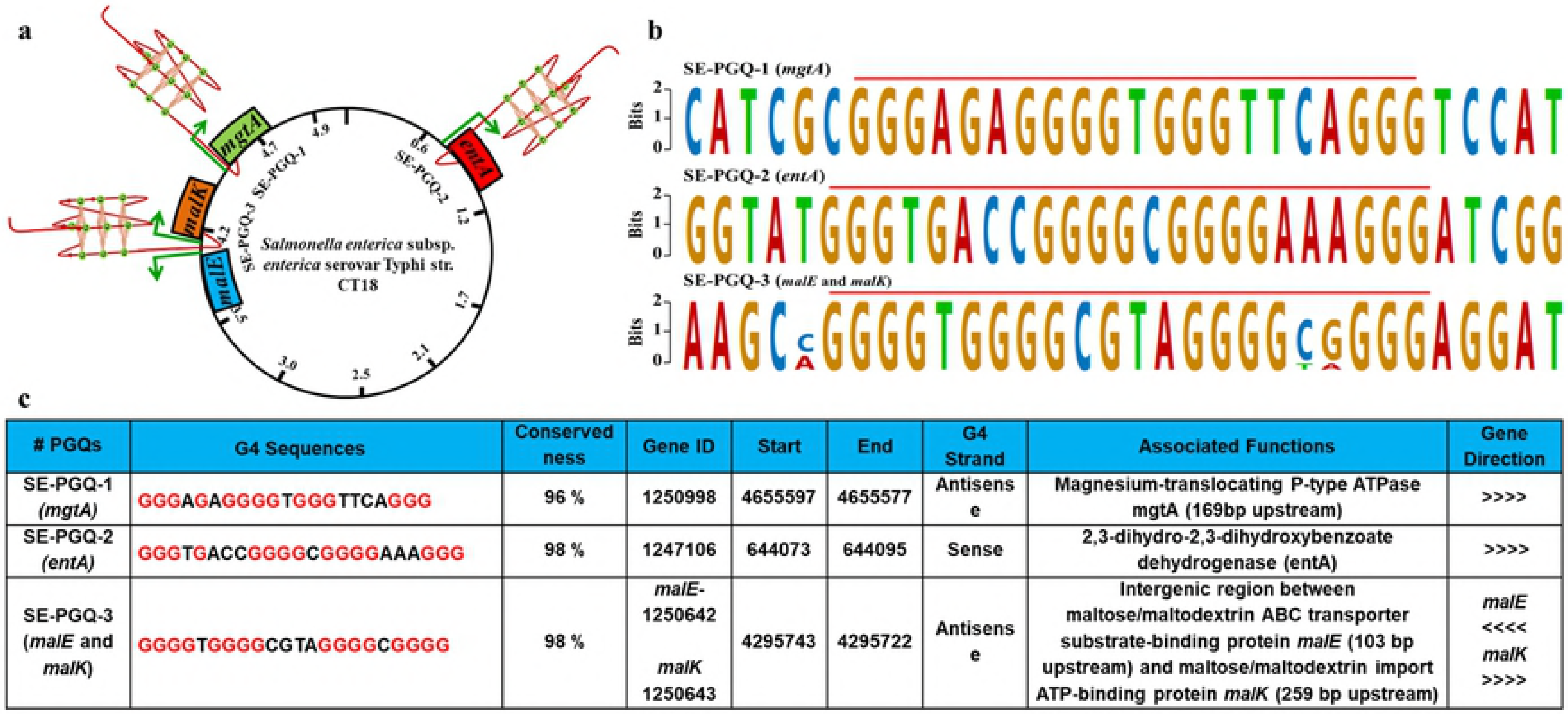
Representation and localization of three highly conserved PGQs. **a)** Diagrammatic representation of SE-PGQ-1 harbored at upstream of *mgtA*, SE-PGQ-2 in the open reading frame of *entA gene*, and SE-PGQ-3 in intergenic regulatory region of *malK* and *malE* in the *Salmonella enterica* genome. **b)** Details of three most conserved PGQs including location, Gene id, Gene locus, G-quadruplex strand and gene direction in the reference genome of *Salmonella enterica* (CT18 strain). **c)** Consensus Sequence of the most conserved PGQs predicted by Glam2 tool of MEME Suite.

As mentioned above, these genes are essential for bacterial survival and virulence inside the host macrophages. Therefore, G-quadruplex motifs found in these genes can be utilized as a potential drug target to develop a promising anti-microbial therapeutics. Moreover, the high conserved-ness of these SE-PGQs, even in the drug-resistant strain would overcome the problem of emergence of drug-resistance in *S. enterica*.

## Result

### *Salmonella enterica* genome harbors three most conserved G-quadruplexes

A comprehensive mining of potential G-quadruplex forming motifs (SE-PGQs) was performed on 412 completely sequenced strains of *S. enterica* (Supplementary Table S1). The bioinformatics analysis observed a total of 109400 PGQs in 412 strains of *S. enterica* (Supplementary File S2). Given that, the similar sequence may correspond to similar structure and evolutionarily conserved function, all the predicted PGQs were further clustered by Unweighted Pair Group Method with Arithmetic Mean clustering method using Clustal Omega tool. The conserved-ness is an essential parameter that makes these PGQ motifs suitable to work as promising drug targets. Therefore, next we examined the conservation of each PGQ clustered using the following equation:

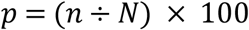

Where *p* is the frequency of occurrence, n = number of strains with specific G4 sequence, and N represents the total number of strains of *S. enterica*. These, conservation analysis revealed 187 PGQ clusters that were observed to possess conversed-ness in more than 90% strains of *S. enterica*. (Supplementary Table S3). G-quadruplex with loop length 1-7 and G tract of ≥3 forms more stable G-quadruplex[50]. Therefore, for the further study, we selected only those PGQs that satisfying the aforesaid criteria of G-quadruplex formation and were listed in Table 1.

**Table 1:**
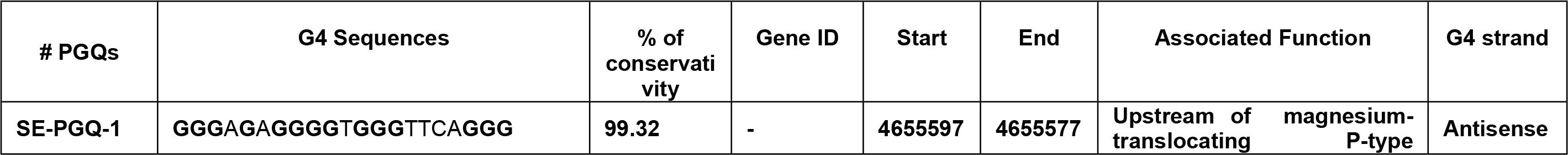
List of PGQs with G-tract ≥3 and loop length 1≤ L ≥7and conserved in 90% *Salmonella enterica* along with their gene IDs, locations, functions, and gene name.

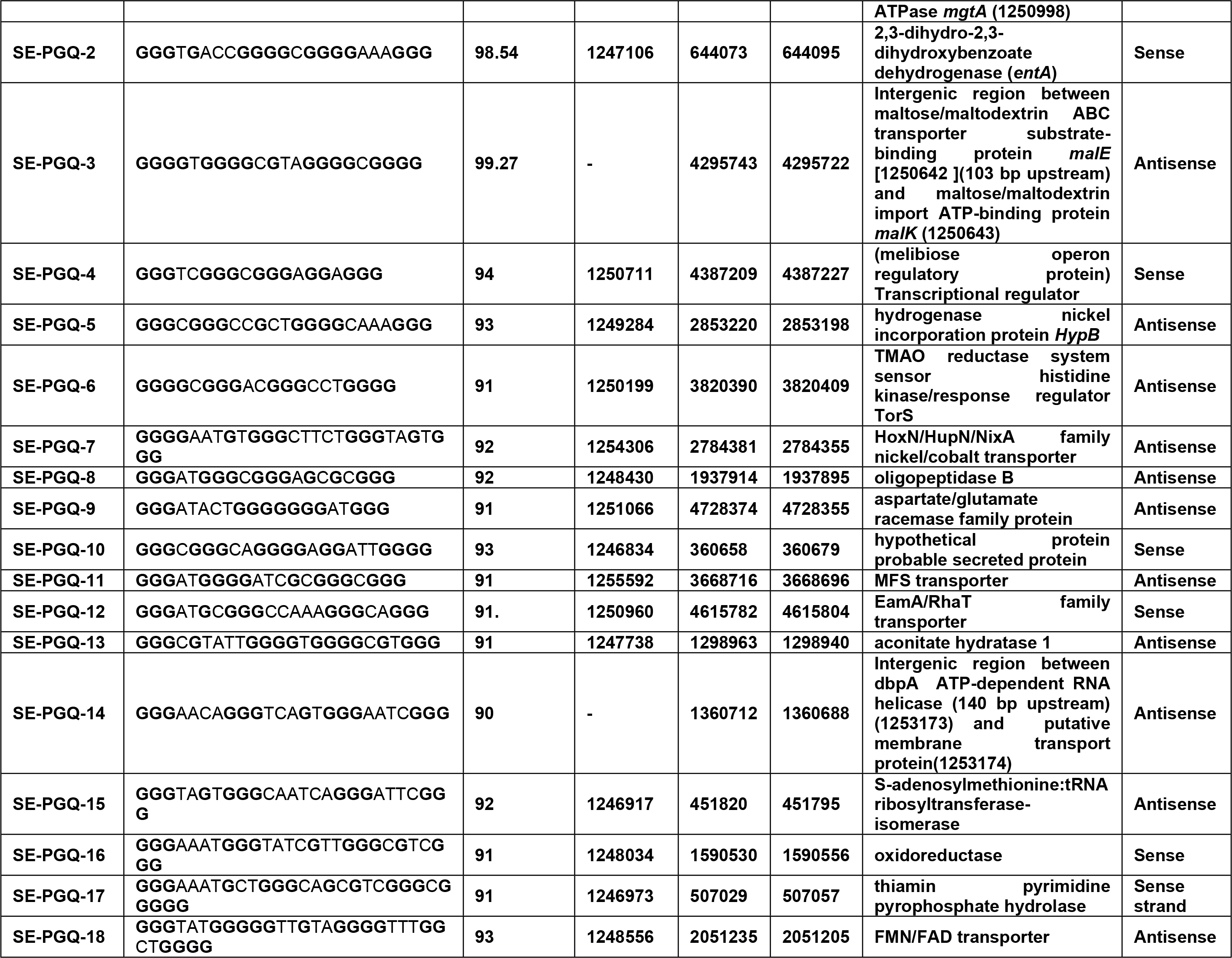

These predictions were further crosschecked with QGRS Mapper and PQSFinder (Supplementary Table S3a and S3b). Interestingly, out of these 18 PGQs clusters, three PGQs (SE-PGQ-1, SE-PGQ-2 and SE-PGQ-3) were found to be conserved in more than 98 % strains of *S. enterica* (S2 File) and present in the four essential genes namely *mgtA*, *entA*, *malK* and *malE* (Fig 3). The consensus sequence depicted the conserved G-residues of SE-PGQ motifs during the evolution process Fig 3b.

### *In vitro* ^1^H NMR analysis affirms the formation of G-quadruplex

NMR spectroscopy is considered as a most reliable technique for confirming the formation of G-quadruplex structure formation by the nucleic acid sequences. Therefore 1D ^1^H NMR spectroscopy was performed to confirm the formation of G-quadruplex conformation by SE-PGQs. The presence of a chemical shift in the range of 10-12 ppm in 1D ^1^H NMR spectra depicts the presence of hoogsteen base pairing in characteristic G-tetrads of G-quadruplex structure whereas, canonical G-C Watson Crick base pairing can be characterized by a chemical shift in the range of 12-14 ppm. All the three SE-PGQs showed an imino proton resonance between 10-12 ppm and clearly affirms the formation of G-quadruplex structure (Fig 4 & S1 Fig).

**Fig. 4.**
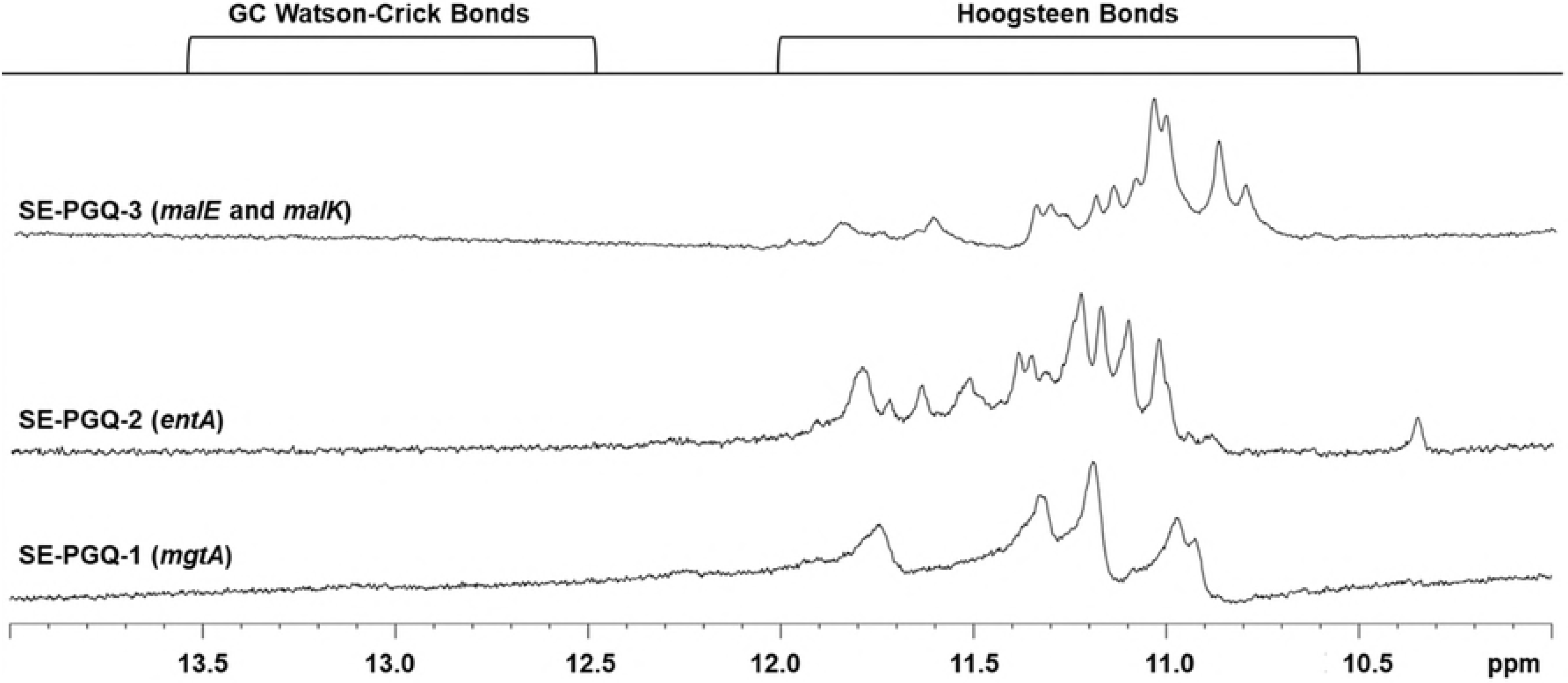
NMR Spectra. 1D ^1^H NMR spectra of SE-PGQ-1(*mgtA*), SE-PGQ-2(*entA*), SE-PGQ-3(*malE* and *malK*) in the presence of K^+^ Buffer.

### Evaluating the topology and stability of the PGQs using Circular Dichroism

Circular dichroism is one of the widely used techniques for analyzing the topology of the G-quadruplex structure. G-quadruplex, depending upon its sequence, loop length, and bound cation, can form either a parallel, antiparallel or hybrid conformation. A positive peak at ~260 nm and a negative peak at ~240 nm signifies for parallel G-quadruplex topology. However, a positive peak at ~290 nm and a negative peak at ~260 nm signifies for anti-parallel G-quadruplex topology whereas, two positive peaks at 260 nm and 290 nm with a negative peak at 240 nm depict the mix or hybrid topology [51]. Different cation affects the stability of the G-quadruplex structure in different extent. The ranking of stabilizing ability of some well studied cations is as follows: K^+^ > Na^+^ > Mg^2+^ > Li^+ [35]^. Therefore, we performed the CD spectroscopy of SE-PGQs in four different cations (K^+^, Na^+^, Li^+^, and Mg^2+^) containing buffers (Fig 5 & S2 Fig).

**Fig. 5.**
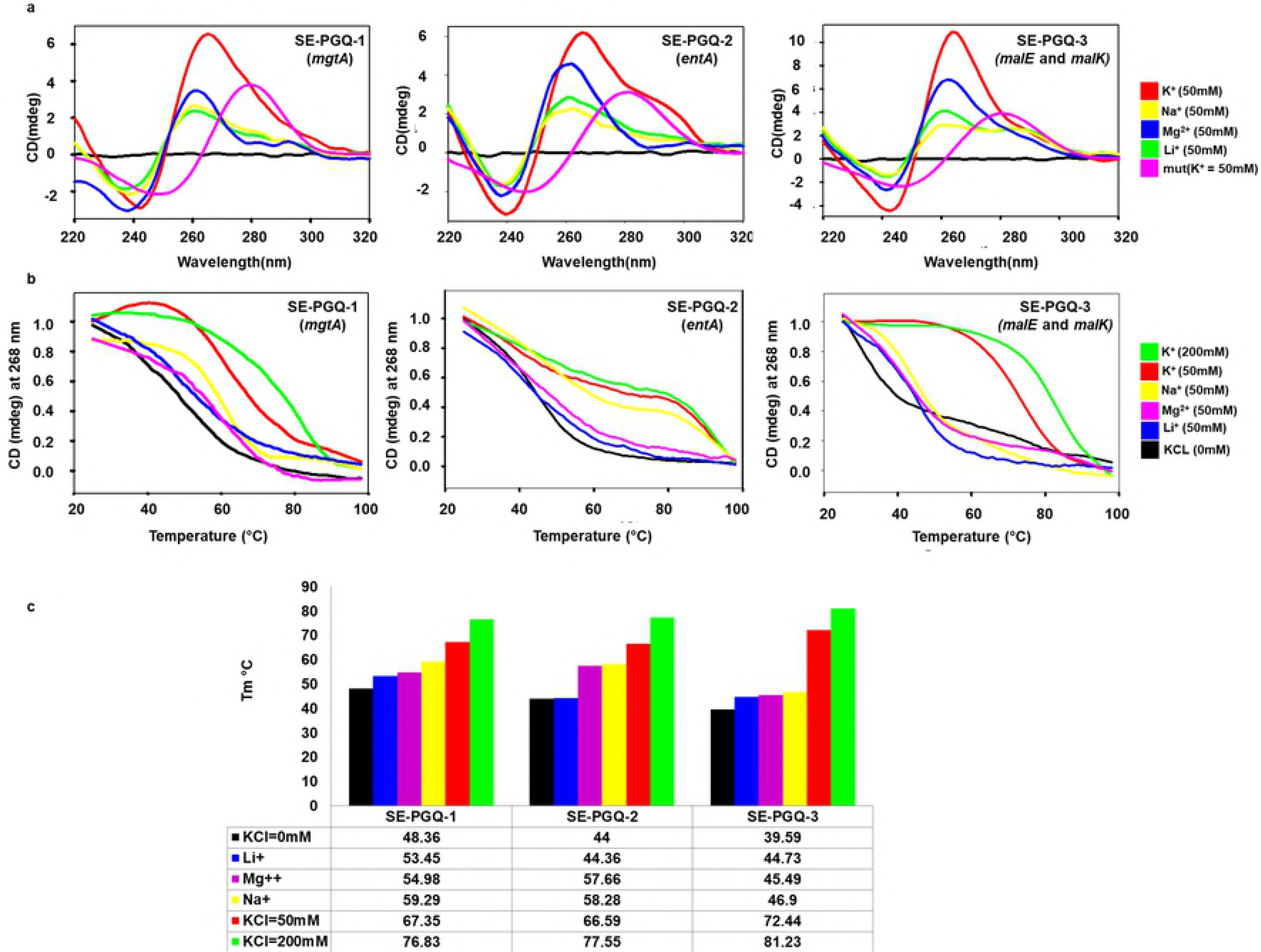
Circular Dichorism Analysis. **a)** Circular Dichroism spectra of the three most conserved PGQs in the presence of Tris-Cl buffer (10 mM) containing either of 50 mM K^+^ (red), 50 mM Na^+^ (green), 50 mM Li^+^(yellow), 50 mM Mg^2+^ (blue) or mut of the same length(Pink). **b)** Melting spectra obtained by Circular Dichroism in different buffers (K^+^, Na^+^, Li^+^ and Mg^2+^) for conserved PGQs predicted in *Salmonella enterica*. In the absence of any buffer(Black), K^+^ 50 mM(Red), K^+^ 200mM (Green), Na^+^ 50 mM (Yellow), Li^+^ 50 mM (Blue)and Mg^2+^ - 50mM (Pink). c) Bar-graph depicting Tm of SE-PGQ-1(*mgtA*), SE-PGQ-2(*entA*), SE-PGQ-3(*malE* and *malK*) in the absence and presence of various cations.

CD spectra analysis revealed the predominant parallel G-quadruplex topology exhibited by SE-PGQ-1 and SE-PGQ-3 in the presence of the K^+^ ion, whereas SE-PGQ-2 showed hybrid G-quadruplex topology in the presence of K^+^ (Fig 5a). As expected, CD spectral scanning performed in the increasing concentration of K^+^ ion showed the maximum molar ellipticity in highest K^+^ ion concentration (S3 Fig). Figure 5b shows the CD melting curve of SE-PGQs in various cation conditions. CD melting analysis revealed the higher stability of the SE-PGQs in the K^+^ ion concentration (Figure 5b & c). To evaluate the significance of G-tracts for their G-quadruplex forming ability, the central Guanine was mutated to Adenine and CD spectra analysis was performed in 50 mM K^+^ ion (S4 Table). Mutants (mut-PGQ-1, mut-PGQ-2, and mut-PGQ-3) failed to show the characteristic CD signal of G-quadruplex i.e. a positive band at 260/290 nm and a negative band at 240 nm suggesting the mutation in G tract disrupted the G-quadruplex formation (Fig 5a).

### Electrophoretic Mobility Shift Assay (EMSA) supports intramolecular conformations of SE-PGQs

Next, Electrophoretic Mobility Shift assay (EMSA) was performed to check the molecularity (inter or intra molecular G-quadruplex) of SE-PGQ in the solution. An intramolecular G-quadruplexes possess a compact topology and migrate faster than their linear counterpart, whereas intermolecular G-quadruplex contains a comparatively wider topology and exhibited slow migration than their linear counterpart[52]. All the three SE-PGQs and positive control (Tel22 DNA G-quadruplex) showed faster mobility than their respective linear counterpart and therefore suggested the formation of intramolecular G-quadruplex by SE-PGQs (Fig 6).

**Fig. 6.**
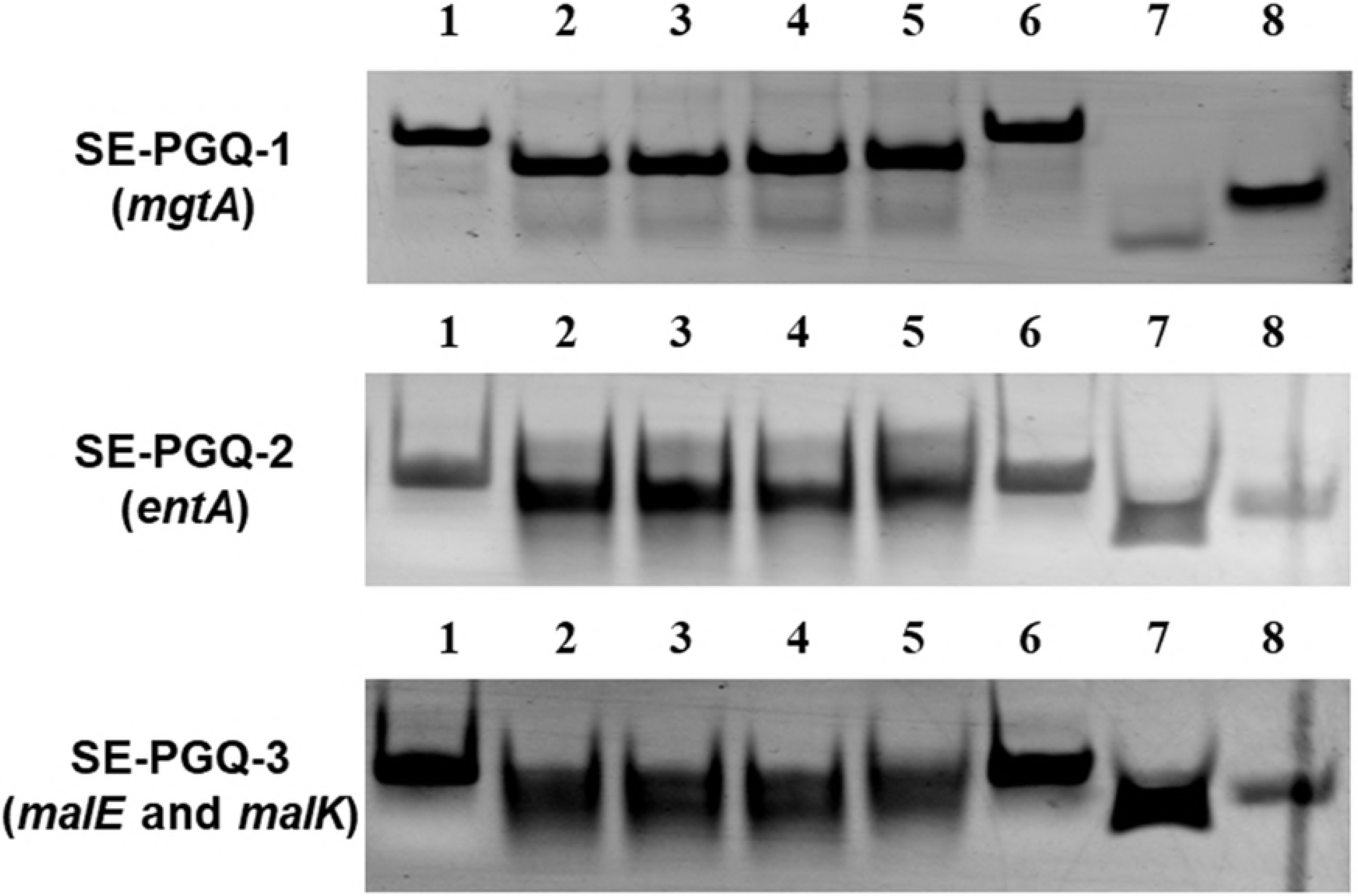
Electrophoretic Mobility Shift assay. Gel images of the SE-PGQ-1(*mgtA*), SE-PGQ-2(*entA*), and SE-PGQ-3(*malE* and *malK*). Lane 1: Mutant (Primer of equal length of the PGQ; Lane 2: PGQ in K^+^ buffer; Lane 3: PGQ in Na^+^ buffer; Lane 4: PGQ in Li+ buffer and Lane 5: PGQ in Mg^2+^ buffer; Lane 6: Mutant (Primer of equal length of the predicted PGQ). Lane7: Positive control (Tel22 G-quadruplex); Lane 8: Negative control (mutant primer of equal length of the Tel22 G-quadruplex sequence)

### 9-amino acridine inhibits *Salmonella enterica growth*

Various small molecules that either stabilized or destabilized the G-quadruplexes conformations are being under investigation for therapeutic intervention of various human pathogenic infection such as BRACO-19, TMPyP4, and several 9-amino acridine derivatives[42]. Previously, 9-amino acridine and its derivatives have been observed for their anti-proliferative properties in cancer cells [53] by binding to the telomeric region [54], the c-Myc gene [55] and c-Kit promoter [56]. Therefore, here we were interested in analyzing the effect of 9-amino acridine on the *S. enterica* growth and performed agar disc diffusion assay and MTT assay. A clear zone of inhibition was observed in the agar plate comparable to ampicillin and penicillin (S4 Fig) that suggested the inhibitory effect of 9-amino acridine on the *S. enterica* growth and an MTT assay observed an IC_50_ value of 10.5 μM (Fig 7).

**Fig. 7.**
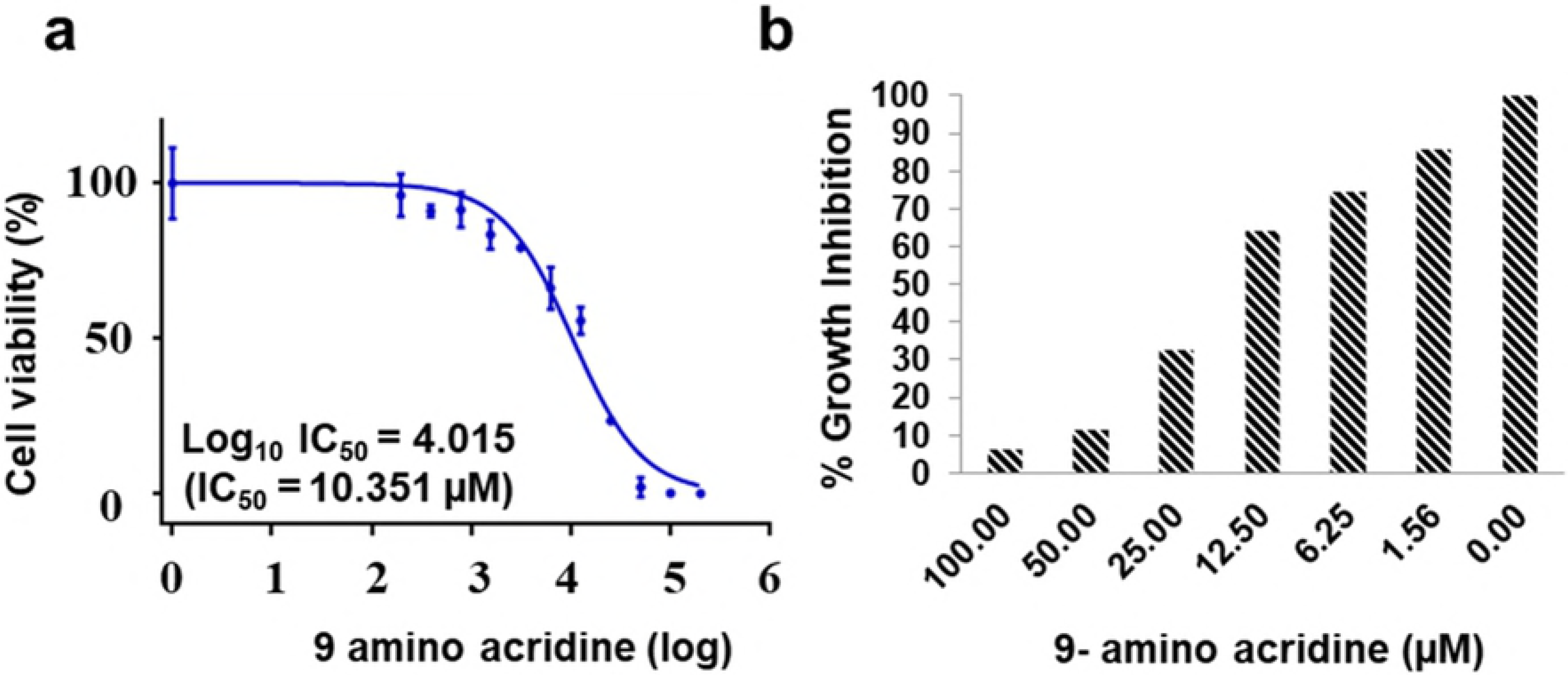
Growth inhibition (MTT) assay of *Salmonella enterica*. **a)** Dose dependent growth curve of *Salmonella enterica* treated with 9-amino acridine. **b)** Growth inhibition of *Salmonella enterica* in the presence of various concentrations of the 9-amino acridine.

### 9-amino acridine stabilized the SE-PGQs and thereby stalls the movement of polymerase

In order to understand the role of SE-PGQs in the cytotoxic effect of 9-amino acridine, binding affinity of 9-amino acridine with these SE-PGQs were analyzed by performing CD Melting studies. An increase in the melting temperature (ΔT_m_~8.5°C) was observed upon addition of 9-amino acridine when compared with alone SE-PGQs. This indicated that 9-amino acridine increased the thermodynamic stability of SE-PGQs (Fig 8). Further, we employed a Taq polymerase PCR stop assay to investigate whether 9-amino acridine complex formation with SE-PGQs, make it possible to stop the movement of polymerase replication machinery or not. In order to investigate this hypothesis, we incubated PCR reaction mixture with 9-amino acridine in a concentration dependent manner and then performed PCR amplification. We observed diminished intensity of bands with increase in concentration of 9-amino acridine, however, in the absence of the 9 amino acridine the band intensity was maximum indicated that the Taq polymerase were able to extend the SE-PGQs motifs. It shows that binding of the 9-amino acridine to the SE-PGQs motif stabilized the G-quadruplex structure and inhibited the movement of replication machinery over the untreated SE-PGQs. On the contrary, when mutant PGQs lacking G-tract were used as a DNA template, 9-amino acridine was not able to bind and thus, could not inhibit the movement of Taq polymerase and produced a PCR product in the reaction(Fig 9).

**Fig. 8.**
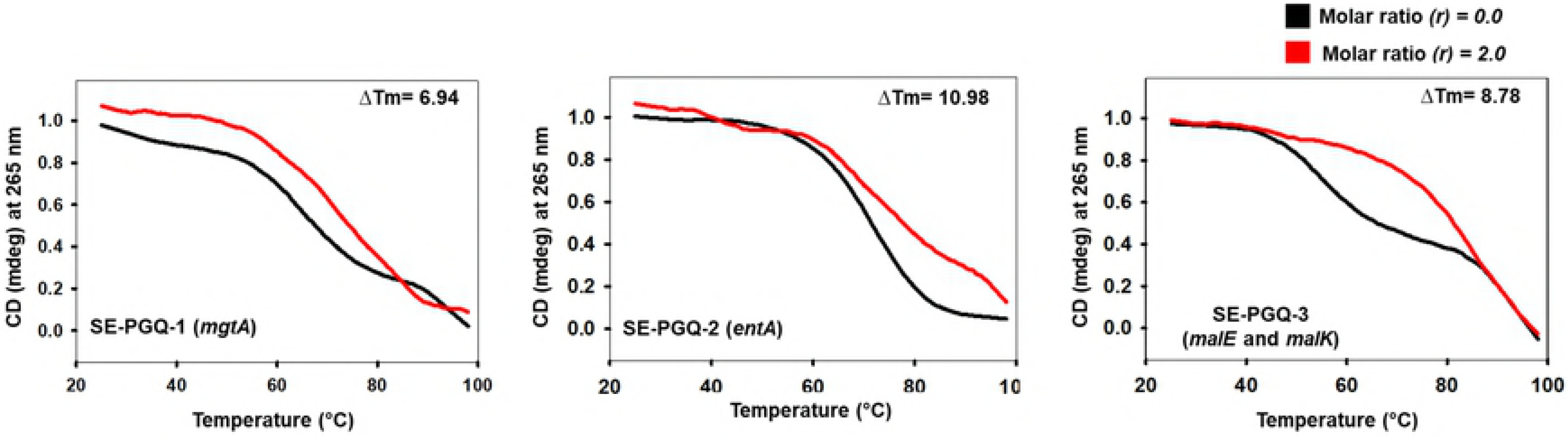
Interaction of 9-amino acridine with SE-PGQs. Circular Dichroism melting curve depicting the change in T_m_ of SE-PGQ-1(*mgtA*), SE-PGQ-2(*entA*), and SE-PGQ-3(*malE* and *malK*) in the presence and absence of 9-amino acridine.

**Fig. 9.**
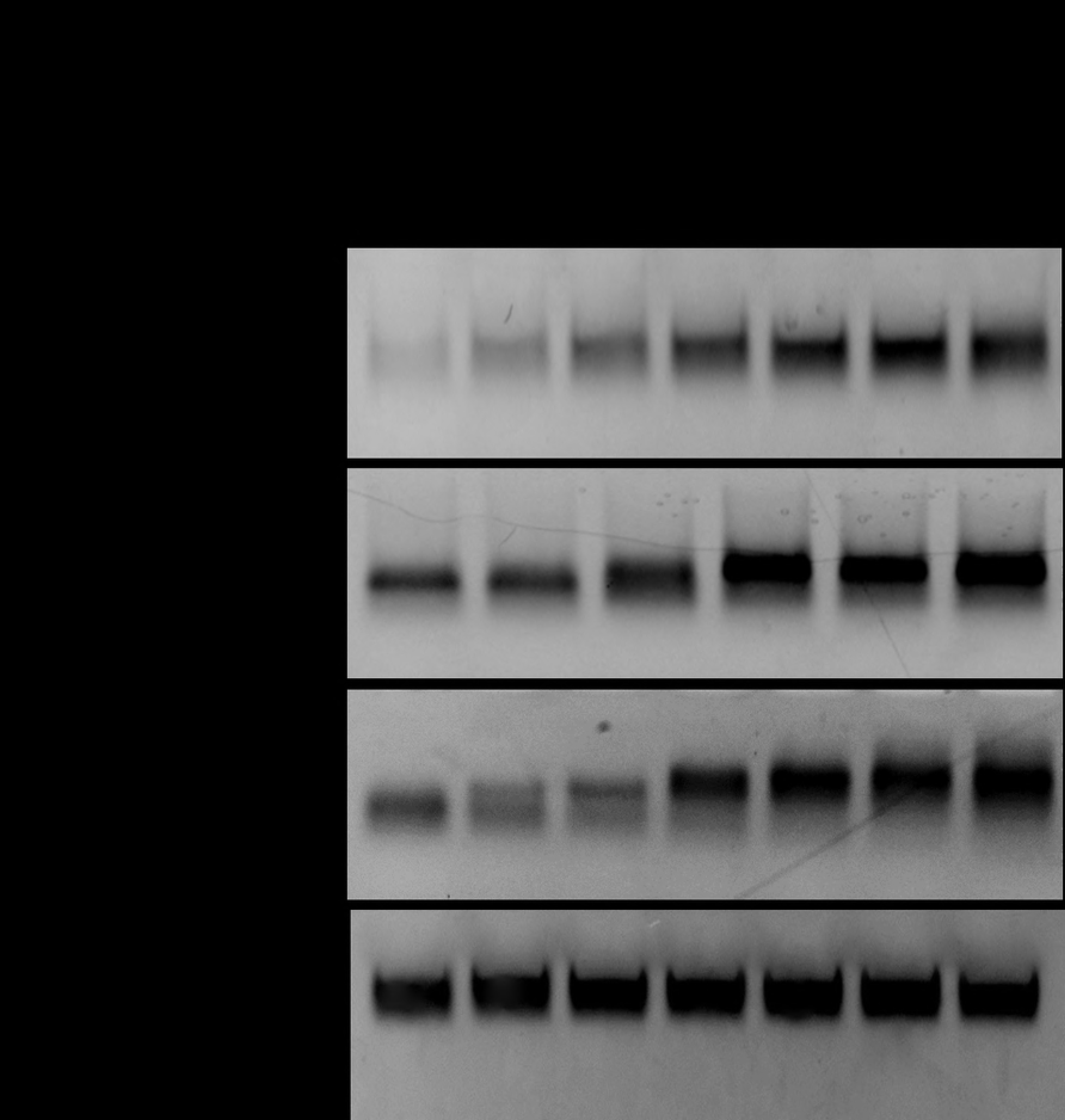
PCR Stop assay. Gel image of Taq Polymerase stop assay for SE-PGQ-1(*mgtA*), SE-PGQ-2(*entA*), and SE-PGQ-3(*malE* and *malK*) and linear DNA with the increasing concentrations of 9-amino acridine.

### 9-amino acridine decreases the transcription rate of the genes harboring PGQs

Further, we performed qRT-PCR assay to check the effect of 9-amino acridine on the expression of the PGQs harboring genes. The gene transcripts of the genes were quantified with respect to 16s rRNA gene and the fold change in the expression of the transcripts was analyzed for the treated cells in comparison to non-treated cell (control culture). The qRT-PCR analysis revealed the presence of the 9 amino acridine reduced the rate of transcription of *mgtA*, *entA* and *malK* by 1.86, 3.03 and 2.94 fold respectively. *malE* gene showed the highest suppression by 7.16 fold decrease in its expression in the presence of 9 amino acridine. In conclusion, all the four genes showed decrease in the expression level, thus strengthening the G4 medicated inhibition mechanism in the their promoter/regulator or open reading frame region (Fig 10)

**Fig. 10.**
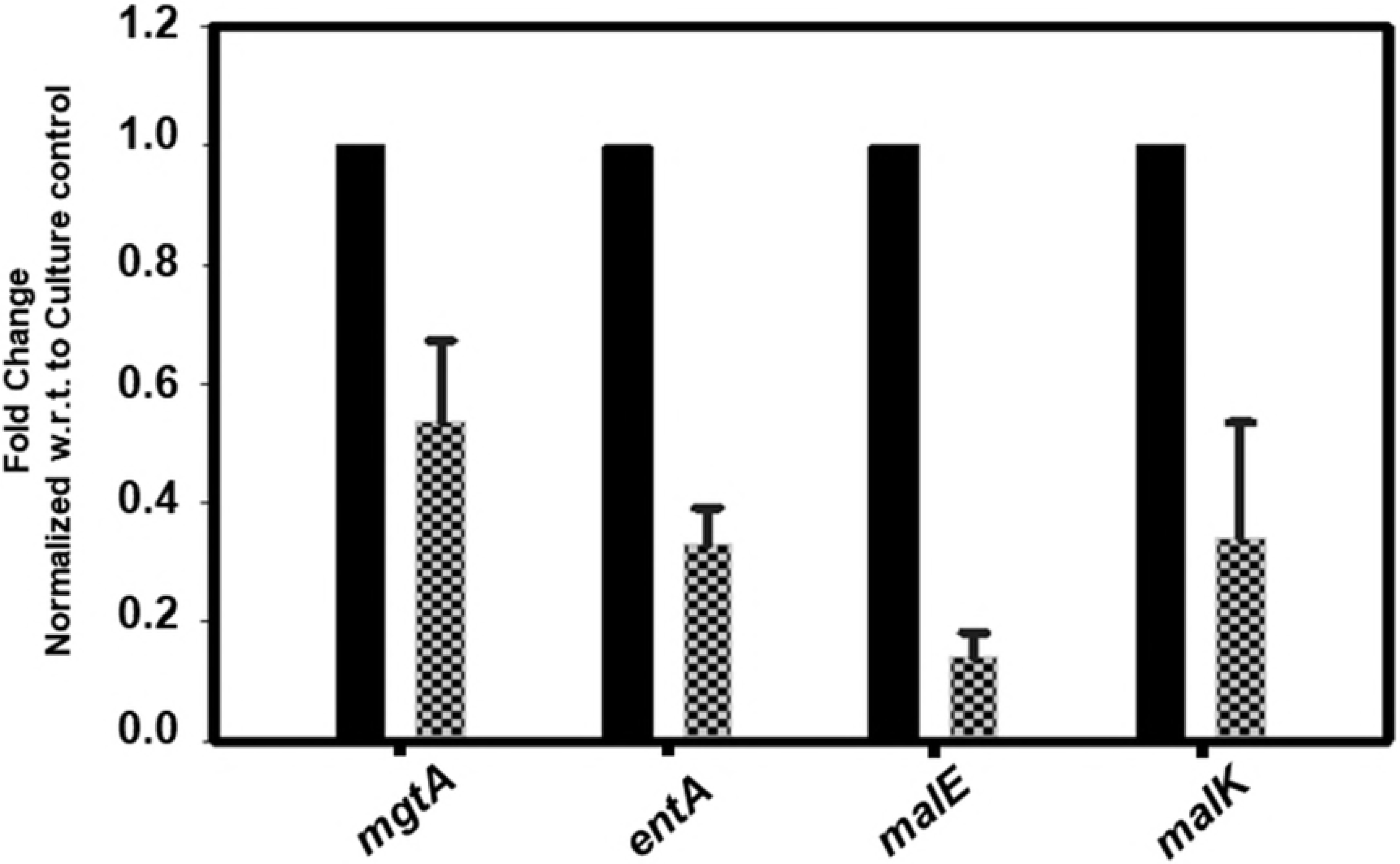
RT-qPCR. The normalized fold change of *mgtA*, *entA*, *malE* and *malK* transcripts in *Salmonella enterica* determined by quantitative PCR in the absence (culture control) and presence of 9-amino acridine(treated).

## Discussion

*Salmonella enterica* is a food-borne pathogenic bacterium and causes a spectrum of diseases including enteric fever (typhoid) and food poisoning in human(2). Global burden of *S. enterica* is rising exponentially due to the emergence of multidrug and extremely resistant strains. Therefore, it becomes essential to develop a novel therapeutic strategy that can work in both drug susceptible and drug-resistant strain infection of this deadly bacterium and it could only be possible if a conserved drug target available in all strains of this bacterium could be targeted. Recently, conserved G-quadruplexes and their binding with small molecule are being extensively investigated as a promising therapeutic approach for combating the various type of human pathogen infection[42, 46]. For example, HIV-1 promoter region possessed a G-quadruplex motif in long terminal repeat (LTR) region of their genome and observed to be critical for its proliferation. BRACO-19 has shown anti-HIV-1 activity by stabilizing G-quadruplex motif present in the LTR region[42]. Similarly, stabilization of G-quadruplex structure present in the core gene of HCV genome by PDP, halts its replication, translation and therefore can be used as potential antihepatitis therapeutics[57]. Pyridostatin is also observed to stabilize the G-quadruplex structure formed in the mRNA of nuclear antigen 1 protein of EBV leading to its translation suppression[58]. Recently, BRACO-19 and quarfloxin have shown inhibitory effect on *Mycobacterium tuberculosis* and *Plasmodium falciparum* by stabilizing G-quadruplexes present in various regions of their genome[40, 59].

Considering the suitability of G-quadruplex structure as a promising drug target in both drug susceptible and drug-resistant strain of pathogens, here we sought to search for G-quadruplex motifs in *S. enterica* strains. The bioinformatics analysis gave a boost to our hypothesis as we received an ample number of potential G-quadruplex motifs in the genome of *S. enterica* strains. Interestingly three G-quadruplex forming motifs were found to be conserved in ≥ 98% strains of *Salmonella enterica* and present in three different locations namely i) SE-PGQ-1 in the regulatory region of *mgtA*, ii) SE-PGQ-2 in open reading frame of *entA* gene. *iii)* and SE-PGQ-3 in the regulatory region shared by maltose operon genes: *malK*, and *malE gene*. 1D ^1^H NMR spectroscopy was used to confirm the formation of G-quadruplex structure by all SE-PGQs whereas EMSA demonstrated the intramolecular G-quadruplex structure formation by SE-PGQs. CD spectroscopy was used to confirm the topology of SE-PGQs in solution. Taken together, the NMR, CD, and EMSA results allow us to conclude that the SE-PGQ-1 and SE-PGQ-3 motifs form intramolecular parallel G-quadruplex structures whereas SE-PGQ-2 form intramolecular mixed type of G-quadruplex structures.

Similar to G-quadruplex motif present in genome of another human pathogen, a G-quadruplex motif present in *mgtA*, *malK*, *malE* and *entA* genes of *S. enterica* strains may also serve as a potential target for developing anti-bacterial therapy. These G-rich targets can also overcome the problem of the drug resistance due to their high conservedness in both drug-susceptible and drug-resistant bacterial strains. Since these PGQs were present in the promoter region of *mgtA*, *malE* and *malK* and the open reading frame of *entA* gene, we explore the stabilizing effect of 9-amino acridine on SE-PGQs. Firstly, CD melting analysis revealed the increased thermos-stability of the SE-PGQs by exhibiting an increase in their melting temperature(T_m_) in the presence of the 9 amino acridine. The observed ΔT_m_ values were observed as 6.94 °C, 10.98 °C and 8.78 °C for SE-PGQ-1, SE-PGQ-2, and SE-PGQ-3 respectively. Further, binding of 9-amino acridine to SE-PGQs reveled the stallation of the movement of DNA replication machinery across G4 motifs. Next, we evaluated the inhibitory effect of 9-amino acridine on *S. enterica* growth; IC_50_ value was calculated to be 10.50 μM. Highly stable G-quadruplexes in the regulatory regions and open reading frame of genes were previously shown to down-regulate the expression of genes [60–62]. The treatment of *S. enterica* cultures with 9-amino acridine led to a decrease in the expression of *entA*, *mgtA*, *malK* and *MalE genes* relative to 16S rRNA, suggesting a G-quadruplex mediated inhibition mechanism is involved in this process. As a schematic model elaborated in Fig 11, G-quadruplex mediated inhibition of *entA*, *mgtA*, *malK* and *MalE genes* is expected to increase the innate immune response of host cell and reduced survival of bacterium inside the host cell. The Inhibited expression of the *entA and mgtA* proteins would make the bacteria unable to response against Reactive oxygen/nitrogen (ROS/RNS)species and reduced their survival inside the host macrophages.

**Fig. 11.**
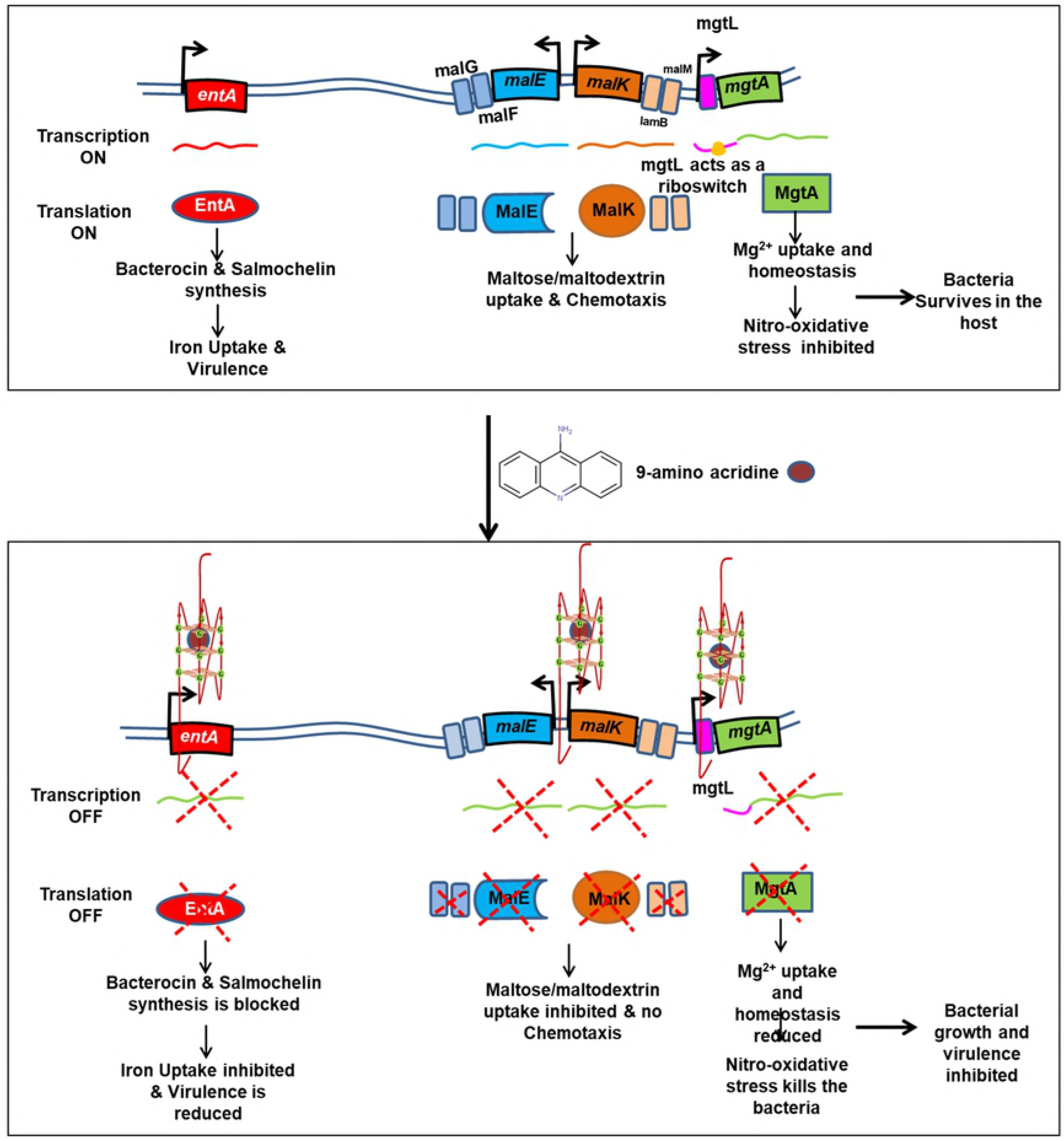
Effect of 9-amino acridine on *mgtA, entA* and maltose operon mediated mechanisms. Schematic representation of G-quadruplex loci and the effect of their stabilization with 9-amino acridine on the survival and virulence of *Salmonella enterica*.

As a concluding remark, the current study shows the presence of stable and highly conserved G-quadruplex structures in essential genes of *Salmonella enterica*. The 9-amino acridine were observed to bind and reduced the expression of these G-quadruplex structure possessing genes and thereby proposed as novel G4 mediated therapeutic approach for combating the infection of *Salmonella enterica* in humans.

## Materials and methods

### Prediction, conserved motif enrichment and functional analysis of G-quadruplex motifs in *Salmonella enterica* strains

Completely sequenced strains of *S. enterica* (S1 Table) were downloaded from National Center for Biotechnology Information (NCBI). These strains were then extensively mined for the potential G-quadruplex motifs in both sense and antisense strand using our previously developed G-quadruplex predictor tool[63]. This prediction tools used the following regular expression.

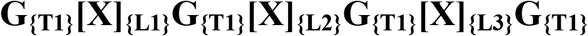

Where T1 represents consecutive tracts of Guanine that can be any number from 2-7, X is any nucleotide (A, T, G, C), L1, L2, L3 represents the variable loop region and can be any number from 1-7. For our prediction, we used G-tracts - 3 or 4 and loop length 1-20 nucleotide [63]. The results were further cross verified by using QGRS Mapper[64] and PGSFinder[65] tools.

To find the conserved PGQ’s that are available in all the strains, multiple sequence alignment (MSA) was performed by using Clustal Omega tool and clustering was done using UPGMA method. Consensus sequences representing the whole G4 sequence with −5 and +5 flanking regions were constructed using the Glam2 tool of MEME Suite[66].

The resultant PGQ clusters were then mapped for their gene location in the genome of the individual *S. enterica* strains using the coordinates extracted from our G4 prediction tool by using Graphics mode of GenBank Database (https://www.ncbi.nlm.nih.gov/nuccore/).

### *Salmonella* genus G4 homolog prediction

In order to check the conservation of predicted PGQs at the *Salmonella* genus level, NCBI nucleotide BLAST was performed by taking each consensus PGQ as a query sequence and *Salmonella bongori* genome sequences as a target(NCBI taxid: 590). The threshold e-value was set as 1e-3 to remove any results by chance.

### Oligonucleotide preparation for CD and ITC analysis

Predicted G4 oligonucleotides sequences were procured from Sigma Aldrich Chemicals Ltd. (St. Louis, MO, USA). 100 μM stock solutions were prepared as per manufacturer’s instructions. Before each set of experiment, oligonucleotides were subjected to re-anneal by heating at 95 °C for 10 minutes and slow cooling at room temperature for 2 hrs. All these oligonucleotides were dissolved in four different Tris-buffer (pH=7.0, 10mM) containing 50 mM of K^+^, Na^+^, Li^+^ and Mg^2+^ separately.

### CD Spectroscopy and melting analysis

CD experiments were performed for each oligonucleotide in different buffer conditions at 25 °C with the scanning rate of 20 nm/min from 220 to 320 nm using Jasco J-185 Spectropolarimeter (Jasco Hachioji, Tokyo, Japan). The instrument was equipped with Peltier Junction temperature controller. Spectra were recorded in a cuvette of 1 mm path length for a final concentration of 20 μM of all oligonucleotides in Tris buffer (10 mM, pH=7.4) containing four different cations *viz*. K^+^, Na^+^, Mg^2+^ and Li+ (50 mM each) in separate experiments. To avoid signal contribution from the buffer, a blank spectrum was recorded before each measurement and subtracted from CD spectrum of the sample prepared in respective buffer.

CD Melting analysis was performed for every PGQs with a temperature range of 25 °C to 98 °C in four different cations (K^+^, Na^+^, Mg^2+^ and Li^+^, 50 mM each) containing buffers. The heating rate was set as 1 °C/min for each melting experiments

### Electrophoretic Mobility Shift assay

Native PAGE was run using 20% polyacrylamide gel in 1X TBE buffer. Each sample was dissolved in Tris buffer(pH=7, 10mM) containing four different cations K^+^, Na^+^, Mg^2+^, Li^+^ (50mM each) separately. For each PGQ, an oligonucleotide of similar length (G mutated with T nucleotide) was taken as a negative control, and standard G-quadruplex (Tel22 DNA) was taken as a positive control. 20 μl of each oligonucleotide sample were loaded, and electrophoresis was performed at 4° C, 90 Voltage in a vertical gel unit system. The gels were visualized by staining with ethidium bromide and analyzed on ImageQuant LAS 4000 gel doc (GE Healthcare Biosciences Ltd., Sweden).).

### One dimensional ^1^H-NMR Spectroscopy

AVANCE 500 MHz BioSpin International AG, Switzerland equipped with a 5 mm broadband inverse probe was used to perform NMR spectroscopic analysis. All the NMR experiments were performed using H_2_O/D_2_O solvent at 9:1 ratio. Temperature of 298K with 20 ppm spectral width and 3 - (Trimethylsilyl) propionic-2, 2, 3, 3-D4 acid sodium salt (TSP) as an internal reference were used. NMR data processing, integration, and analysis were done by using Topspin 1.3 software.

### Bacterial strain culture and growth conditions

The S. ser. Typhimurium strain ATCC 14028 was procured from HiMedia and streaked on Nutrient Agar (HiMedia). A single colony was inoculated in the in Nutrient Broth (HiMedia) and kept overnight at 37°C and 220 rpm in incubator shaker.

### Growth inhibition assay - Disk Diffusion and MTT

Disk diffusion method was employed to identify the susceptibility of bacteria to 9-amino acridine. For the antimicrobial activity of 9-amino acridine to *S. enterica*, the overnight growth culture was spread onto the Nutrient Agar plate. Whatman paper disks soaked with 9-amino acridine solution (in DMSO) containing 20 μg, 10 μg, and 5 μg placed onto the plates spread with *S. enterica*. Penicillin and Streptomycin (HiMedia) standard disk of 10 μg were used as positive control while the solvent, DMSO was used as a negative control. The plates were incubated at 37°C for 16 hours, followed by measurement of the zone of inhibition [67].

MTT assay was performed for cytotoxic analysis of 9-amino acridine on *S. enterica*. 50 μl of the overnight grown culture of *S. enterica*, was inoculated in 5 ml Nutrient Broth (NB) at 220 rpm, 37°C and allowed to grow till the O.D_600_ =0.5. After that, 50 μl was transferred in fresh 5 ml NB tube and 100 μl was transferred in each well of 96 well plate. Dilution was prepared from the stock solution (200 μM) of 9-amino acridine of the following concentrations 100 μM - 0.09 μM and added to the respective wells, last well served as blank (without 9-amino acridine). The plates were kept at 37°C, 220 rpm for 3 hr. Afterward, 10 μl of MTT (5 mg/mL) was added to each well and incubated for 3 hr. Finally, 20 μl of DMSO was added in each well to dissolve formazan crystal, and the plate was examined using microplate reader (BioTek) at 590 nm[68].

### PCR Stop Assay

Templates and Primers were procured from Sigma-Aldrich Chemicals Ltd. (St. Louis, MO, USA) (Table S5a). The reaction was performed in the master mix consisting of 1X PCR buffer, 0.33 mM dNTPs, 2 templated, 2 μM reverse primer, 2.5 units Taq DNA polymerase (Sigma-Aldrich Chemicals Ltd. St. Louis, MO, USA) and a dose titration of 9-amino acridine. The following thermal cycling conditions were used: initial denaturation at 95 °C for 5 min, followed by 25 cycles of 95 °C for 30 s, 50 °C for 30 s, 72 °C for 0.5 minute and finally held at 4 °C. The 6X DNA loading dye was added to the amplified products and resolved in 3% agarose gel. Ethidium bromide was used for staining and gel images were analyzed using ImageQuant LAS 4000 (GE Healthcare, Biosciences Ltd.,Sweden).

### Total RNA isolation and cDNA synthesis

*S. enterica* was grown overnight at 37°C, 220 rpm in 5 ml of test tube containing NB. Two flasks of 500 ml containing 50 ml of NB each were inoculated with 1% inoculum from overnight culture. Both the flasks of untreated and treated cells were allowed to grow according to previously mentioned condition till the O.D_600_ nm reached 0.5. At O.D_600_ = 0.5, 10.35 μM of 9-amino acridine were added to MIC flask. For control culture DMSO was added as 9-amino acridine was dissolved in it. All the flasks were kept at 37°C, 220 rpm for 45 min. Subsequently, samples were centrifuged at 12000 rpm and immediately preceded for RNA isolation. *S. enterica* culture prior to RNA isolation was treated with RNA protect reagent (Qiagen, USA) to stabilize RNA and prevent it from degradation. Treated and untreated *Salmonella* cultures were used for total RNA isolation. RNA isolation for all samples was carried out using TRIZOL reagent (Invitrogen) according to manufacturer’s instructions. After RNA isolation, its concentration and purity was measured using NanoDrop (Thermoscientific) as ng/ul and A260/A280, respectively. Subsequently, all the RNA samples were treated with RNase-free DNaseI (Invitrogen) as mentioned by the manufacturer. Finally, 5 μg of total RNA from each sample were used for cDNA synthesis. cDNA synthesis was performed in a 20 μl reaction by using Invitrogen Superscript IV kit (Cat # 18091050) according to the manufactures instruction.

### Gene Expression Profiling Using Real-Time quantitative PCR

Quantitative Real-Time PCR was used to elucidate the variation in gene expression profiling of *Salmonella* treated with 9-amino acridine. All the aliquots of cDNA from untreated (control) and treated (9-amino acridine) were used in real time PCR. qPCR was carried out in PCR master reaction mix containing 1x PowerUp™ SYBR^®^ Green Master Mix (Applied Biosystem, USA), 0.5 μM of each primer and 11 μl of cDNA sample in a final reaction volume of 25 μl in a 96 well PCR plate in Step One Plus (Applied Biosystem, USA) Thermal Cycler. All the samples for real time were analyzed in three dilutions and duplicates, normalization was done with respect to 16s rRNA (housekeeping gene). The proportionate change in genes expression was assessed by change in expression of target gene in treated as compared to control. List of primers used in real time PCR and thermo-cycling conditions are mentioned in Supplementary Table S5b and Supplementary Table S5c, respectively. Briefly, the thermocycle used in qRT-PCR was 94°C for 2 min, subsequently 40 cycles of 94°C for 15 sec and 57°C for 1 min.

## AVAILABILITY

Putative G-quadruplex prediction tool is an open web-based tool to predict putative G-quadruplex-forming sequences in different nucleotide stretches. (http://bsbe.iiti.ac.in/bsbe/ipdb/pattern2.php)

## Authors Contribution

Data conceptualization and methodology was performed by A.K. Bioinformatics analysis was done by S.K.M., N.J., and U.S. Biophysical analysis was performed by S.K.M., N.J., U.S. S.K.M. N.J. and A.J. performed MTT assay. A.J., S.K.M. and N.J. performed RT-PCR under the supervision of P.K. and A.K. N.J., S.K.M. and A.T. wrote the manuscript. A.K. T.K.S. and P.K. did the review and editing.

## Acknowledgment

The authors are thankful to SIC Facility at IIT Indore for NMR and CD experiments.

## Supplementary Files captions and Legends

### S1 File. G-quadruplex prediction

List of G-quadruplexes predicted in 412 strains of Salmonella enterica with G-tract 3 and 4 with loop length 0 to 20 bp.(Excel file).

### S2 File. Cluster of highly conserved SE-PGQs in the Salmonella enterica strains

Position of SE-PGQ-1(mgtA), SE-PGQ-2(entA) and SE-PGQ-3(malE and malK) in the strains of *Salmonella enterica* genome. (Excel file)

### S1 Fig. NMR Spectra of 15 PGQs conserved in more than 90% strains of *Salmonella enterica*

1D 1H NMR Spectra of PGQs conserved in ≥90 %(SE-PGQ-4 to SE-PGQ-18) in K^+^ buffer.(TIFF file)

### S2 Fig. CD Spectra of 15 PGQs conserved in more than 90% strains of *Salmonella enterica*

Circular Dichroism spectra of SE-PGQ-4 to SE-PGQ-18 in Tris-Cl buffer (10 mM) containing either 50 mM KCl (red), 50 mM NaCl (green), 50 mM LiCl (yellow) or 50 mM MgCl2 (blue). (TIFF file)

### S3 Fig. CD Spectra with increasing concentration of K^+^ ion

Circular Dichroism spectra of the most conserved PGQs in the absence of any cation or in the presence of 50 mM K^+^ (Green), 100 mM K^+^ (Yellow), 150 mM K^+^ (Blue), or 200 mM K^+^ (Pink). (TIFF file)

### S4 Fig. Agar Disc diffusion assay

Zone of inhibition obtained by agar disc diffusion assay for 20, 10 and 5μg of 9-amino acridine against *Salmonella enterica*. (**TIFF file**)

### S1 Table. Salmonella enterica strains used in the analysis

List of *Salmonella enterica* strains taken into consideration for our analysis with their genome IDs.(**Word file**)

### S2 Table. Conserved G-quadruplex in Salmonella enterica

Complete list of G-quadruplexes in *Salmonella enterica* conserved in more than 90% of strains with G-tract 3/4 and loop length 0 to 20 bp. (**Word file**)

### S3 Table: Conserved G-quadruplex prediction using QGRSMapper and PQSFinder

**a)**. G4 prediction of conserved PGQ’s using QGRS mapper tool. b). G4 prediction of conserved PGQ’s using PGSFinder tool. (**Word file**)

### S4 Table. Mutants for CD analysis

List of mutant sequence used for Circular Dichroism analysis. (Mutated A is highlighted with red) (**Word file**)

### S5 Table. Primer List

**a)** List of primers used for Taq polymenrase stop assay. **b)** List of primers used in RT-PCR. **c)** List of parameters used for thermal cycling conditions used for RT-PCR assay. (**Word file**)

